# IFIT3 and IFIT2/3 promote IFIT1-mediated translation inhibition by enhancing binding to non-self RNA

**DOI:** 10.1101/261776

**Authors:** Renata C Fleith, Harriet V Mears, Edward Emmott, Stephen C Graham, Daniel S Mansur, Trevor R Sweeney

## Abstract

Interferon-induced proteins with tetratricopeptide repeats (IFITs) are highly expressed during the cell-intrinsic immune response to viral infection. IFIT1 inhibits translation by binding directly to the 5′ end of foreign RNAs, particularly those with non-self cap structures, precluding the recruitment of the cap-binding eukaryotic translation initiation factor 4F and subsequent 40S recruitment. Interaction of different IFIT family members is well described, but little is known of the molecular basis of IFIT association or its impact on function. Here, we reconstituted different complexes of IFIT1, IFIT2 and IFIT3 *in vitro*, which enabled us to reveal critical aspects of IFIT complex assembly. IFIT1 interacts rapidly and strongly with IFIT3 forming a stable heterotetramer. IFIT2 and IFIT3 homodimers dissociate to form a more stable heterodimer that associates with IFIT1, forming an IFIT1:IFIT2:IFIT3 trimer. Site-directed mutagenesis revealed a C-terminal ‘YxxxL’ motif in IFIT1 that mediates its association with IFIT3. Using various reporter mRNAs, we demonstrate for the first time that IFIT3 stabilises IFIT1 binding to cap0-mRNA and enhances its translation inhibition activity. Disrupting the binding interface between IFIT1 and IFIT3 abrogated this enhancement. This work reveals molecular aspects of IFIT assembly and provides an important ‘missing link’ between IFIT interaction and function.

## Introduction

The host innate immune response provides a first line defence against invading pathogens. Following infection, pathogen recognition receptors (PRRs) sense non-self, pathogen-associated molecular patterns (PAMP_s_) triggering signalling pathways that activate an immune response (reviewed in 1). Detection of viral signatures by PRRs, such as RIG I-like receptor sensing of double stranded RNA, induces production of Type I and Type III interferon (IFN). Through binding of cell surface IFN receptors and subsequent activation of the JAK-STAT pathway, IFN activates the transcription of hundreds of IFN-stimulated genes (ISGs) (2), many with known antiviral properties, priming neighbouring cells to restrict viral spread.

The **<u>*i*</u>**nter**<u>*f*</u>**eron **<u>*i*</u>**nduced protein with **<u>*t*</u>**etratricopeptide repeats (IFIT) protein family, present in all vertebrates, include some of the most highly expressed ISGs. IFIT1, IFIT2, IFIT3 and IFIT5 contain multiple IFN-stimulated response elements (ISREs) in their promoters (reviewed in 3) and are also induced directly by the IFN-regulatory factor 3 (4), downstream of initial PRR activation. In contrast, the promoter of the poorly characterised human IFIT1B lacks ISREs. Different species have varying complements of IFITs, but most mammals possess IFIT1, IFIT1B, IFIT2, IFIT3 and IFIT5 (5). Phylogenetic analysis revealed that rodents, including mice, have lost IFIT1 and instead Ifit1b1 is IFN responsive (6). In mice Ifit1b1 has undergone duplication twice (Ifit1b2 and Ifit1b3) and Ifit3 once (Ifit3b) (6). IFITs are composed of sequential tetratricopeptide repeat (TPR) motifs that form globular N- and C-terminal domains joined by a linker of variable flexibility (7–10).

TPR motifs are frequently involved in protein-protein interactions and are commonly found in scaffolding proteins (11). The crystal structures of IFIT1 and IFIT5 revealed a positively charged pocket formed in the groove between the N and C domains that interacts with single-stranded RNA (8–10). IFIT1 RNA binding activity was first reported by Pichlmair *et al.* (12) who identified proteins in lysates from IFN stimulated cells that interacted with 5’-ppp RNAs. Subsequently, we and others demonstrated that IFIT1 tightly binds capped RNAs lacking methylation on the first cap-proximal nucleotide (cap0) with low nanomolar affinity (13–15). While similar positively charged tunnels in IFIT1 and IFIT5 interact with the phosphate backbone of bound RNAs (8, 9, 13), only IFIT1 possesses a large hydrophobic cavity at the rear of the tunnel that can accommodate the cap structure (9). These findings support a model whereby IFIT1 out-competes eukaryotic initiation factor (eIF) 4E/4F for binding to cap0-mRNAs, thereby inhibiting their translation. As host mRNAs are generally methylated on the first or first and second bases (cap1 and cap2, respectively), this selectivity offers a mechanism of recognising and blocking translation of non-self RNAs.

Viruses have evolved numerous mechanisms to circumvent detection and inhibition by IFIT1. For example, members of the *Flaviviridae* and *Coronaviridae* families that replicate in the cytoplasm and rely on cap-dependent translation of their single-stranded positive-sense RNA genomes, encode their own 2’-*O*-methyltransferases. Disruption of methyltransferase activity increases susceptibility of the flaviviruses West Nile virus, japanese encephalitis virus and dengue virus (15–19) and the coronaviruses murine hepatitis virus and severe acute respiratory syndrome virus (20, 21) to IFIT1 restriction. Interestingly, enzymatically 2’-*O*-methylated, capped mRNAs from parainfluenza virus 5 display differential translational sensitivity to IFIT1 *in vitro*, while wild type middle east respiratory syndrome coronavirus (MERS) replication was enhanced upon IFIT1 depletion suggesting that factors other than 5’ end methylation can influence IFIT1 recognition (22, 23). By contrast, alphaviruses, such as the emerging human pathogen chikungunya virus, also rely on cap-dependent translation but lack a virally encoded 2’-*O*-methyltranferase, thus possessing viral mRNAs with a cap0 structure at the 5’ end (24). Recent evidence suggests that stable secondary structure at the 5’ end of alphaviral genomes protects the viral RNAs from IFIT1 restriction (25, 26). IFIT1 may also affect translation through interaction with eIF3 (27), while its direct binding to the viral E1 protein restricts human papilloma virus replication (28).

An intriguing feature of IFITs is their propensity to homo and heterooligomerise. Using pull-down experiments of differentially tagged IFITs, Stawowczyk *et al.* (29) demonstrated that IFIT2 could interact both with itself and with IFIT1 and IFIT3 in HeLa cells. In the same study, an IFIT1, IFIT2 and IFIT3 containing complex in HeLa cytoplasmic lysates was also reported to migrate between 150-200 kDa when analysed by glycerol gradient sedimentation. Deletion analysis identified the first four TPRs of IFIT2 as being important for interaction with IFIT3 while the TPR(s) of IFIT2 that promote interaction with IFIT1 could not be elucidated (29). IFIT1 was later reported to interact with IFIT2 or IFIT3 by size exclusion chromatography (SEC) (12). The crystal structure of IFIT2 revealed it forms a stable, domain-swapped dimer, with TPRs of the N-terminal domain exchanged between each monomer (7). Using native gel electrophoresis, we previously demonstrated IFIT1 and IFIT3 could homooligomerise (13), while a recently-reported crystal structure of IFIT1 identified a motif in the C terminus responsible for IFIT1 dimerisation (BioRxiv: https://doi.org/10.1101/152850).

Despite considerable evidence for IFIT oligomerisation, little is known about how different IFITs interact and what impact this interaction has on function. To address this, here we have reconstituted different IFIT complexes from individually purified proteins, determined the oligomeric state of each and reveal for the first time that interaction with IFIT3 or a heterocomplex of IFIT2 and IFIT3 enhances the cap0 RNA binding and translation inhibition activity of IFIT1. We also demonstrate *in vitro* that although IFIT2 and IFIT3 individually form stable homodimers, the IFIT2:IFIT3 dimer complex is the energetically more stable species. Our results provide a critical missing link between IFIT oligomerisation and function, and present a mechanistic framework for understanding the role of IFITs in the host immune response.

### Material and Methods

#### Plasmids

For mammalian cell expression, sequences for human IFIT1 (BC007091.1) and IFIT5 (BC025786.1) were PCR amplified to include a 5′ Kozak sequence, 3′ FLAG tag and 5′ BamHI and 3′ XhoI sites, to facilitate cloning into pCDNA3.1. The plasmid for expression of IFIT1 in *Escherichia coli* was previously described (10) and was used as a template for site directed mutagenesis to generate the mutant IFIT1 expression vectors. Sequences for IFIT2 (NM_001547.4) and IFIT3 (NM_001549.5) were PCR amplified to contain 5′ NdeI and 3′ XhoI restriction sites for cloning into pET28b (Novagen) producing a full-length protein with thrombin cleavable N-terminal 6-His tag. For reporter RNA transcription, the firefly luciferase reporter gene (Fluc) was PCR amplified using primers containing the 5′ UTR and 3′ UTR sequences of human β-globin (NM_000518.4), including a 5′ T7 promotor and EcoRI and PstI sites to facilitate cloning into pUC57. pUC57-ZIKV-Fluc was previously described (30).

#### Protein expression and purification

Recombinant IFITs were expressed in Rosetta 2 (DE3) pLysS (Novagen). Cells were grown to an OD_600_ of approximately 1 in 2x TY media at 37 °C. Expression was induced by adding 1 mM isopropyl β-D-1-thiogalactopyranoside. The induced culture was incubated at 22 °C for 16 hours. Cells were harvested and lysed in a buffer containing 20 mM Tris-Cl pH 7.5, 400 mM KCl, 5 % glycerol, 1 mM DTT and 0.5 mM phenylmethylsulfonyl fluoride and 0.5 mg/ml lysozyme (from hen egg). IFITs were isolated by affinity chromatography on Ni-NTA Agarose beads (Qiagen). IFIT1 and IFIT1 mutants were additionally purified by FPLC on MonoQ (Q buffer: 20 mM Tris-Cl pH 7.5, 5 % glycerol, 1 mM DTT and 100-500 mM KCl), followed by MonoS 5/50 GL (S buffer: 30 mM HEPES pH 7.5, 5 % glycerol, 1 mM DTT and 100-500 mM KCl). IFIT2 and IFIT3 were treated with thrombin (from bovine plasma).

IFIT2 and IFIT3 were further purified on MonoQ 5/50 (Q buffer), followed by size exclusion chromatography (SEC) on Superdex 200 increase 10/300 GL or HiLoad 16/600 Superdex 200 pg columns (SEC buffer: 20 mM Tris-Cl pH 7.5, 150 mM KCl and 1 mM DTT). All FPLC columns are from GE Healthcare and all buffer reagents and enzymes were from Sigma.

#### Western blotting

Proteins were resolved by 12.5 % SDS-PAGE and transferred to 0.45 μm nitrocellulose membrane. For immunoprecipitations, membranes were probed with anti-FLAG M2-Peroxidase (A8592, Sigma), detected by chemiluminescence using Westar Supernova substrate (Cyanagen) and visualised on Super RX-N film (Fugifilm). As a loading control, membranes were probed with anti-GAPDH (AM4300, ThermoFisher), detected using IRDye 800CW Goat anti-Mouse and visualised on an Odyssey CLx Imaging System (Li-Cor). To normalise recombinant IFIT proteins and complexes, membranes were probed with anti-penta-His (34660, Qiagen) and quantified using ImageJ.

#### IFIT complex assembly

All complexes were assembled in SEC buffer.

**IFIT1:IFIT2** and **IFIT1:IFIT3-** 1 mg/mL of each protein was mixed and incubated for one hour at 4 or 30 °C. **IFIT2:IFIT3-** 0.18 mg/mL of each protein was mixed and incubated for one hour at 37 °C and concentrated to 2 mg/mL. **IFIT1:IFIT2:IFIT3 trimer-** 0.4 mg/mL of purified IFIT2:IFIT3 complex and 0.2 mg/mL of IFIT1 were incubated for 1 hour at 30 °C. The complex was concentrated to 3 mg/mL. **IFIT1:IFIT2:IFIT3 tetramer-** 0.4 mg/mL of purified IFIT2:IFIT3 complex and 0.4 mg/mL of IFIT1 were incubated for 1 hour at 30 °C. The complex was concentrated to 4 mg/mL. Complexes were concentrated using Amicon Ultra 0.5 mL 10 kDa molecular weight cut off filters (Millipore).

#### SEC analysis of mutant IFIT1 and IFIT3 complexes

Wild type or mutant IFIT1 was combined with IFIT3 as described in *IFIT complex assembly,* above. 150 µL of each assembly reaction was injected on to a Superdex 200 Increase 10/300 GL column, at 0.3 mL/min flow rate and UV280 readings were monitored. Peak fractions were analysed by SDS-PAGE and Coomassie staining.

#### SEC - Multi-Angle Light Scatter (SEC-MALS)

Proteins/complexes were injected (100 µL at the concentration described in *IFIT complex assembly, above*) onto an analytical Superdex 200 Increase 10/300 gel filtration column. MALS analysis was performed at room temperature, by inline measurement of static light scattering (DAWN 8+, Wyatt Technology), differential refractive index (Optilab T-rEX, Wyatt Technology), and 280 nm absorbance (Agilent 1260 UV, Agilent Technologies) following SEC at a flow rate of 0.4 mL/min. Molecular masses were calculated using the ASTRA6 software package (Wyatt Technology).

#### Differential scanning fluorimetry

Differential scanning fluorimetry experiments to determine the melting temperature (T_M_) of IFIT2, IFIT3 and IFIT2:IFIT3 were performed using a Viia7 Real-Time PCR system (Applied Biosystems). In an optical 96-Well Reaction Plate (Applied Biosystems 4366932), 1:500 Protein Thermal Shift dye (Life Technologies, 4461146) was mixed with 0.1 mg/mL protein in a final buffer composition of 20 mM HEPES pH 7.5, 150 mM KOAc, 2.5 mM MgOAc, 5% glycerol, and 1 mM DTT in a final volume of 20 μL. Emission from quadruplicate samples was measured at 623 nm while ramping from 25 – 95 °C stepwise at a rate of 1 °C per 20 seconds. To determine T_m_, data were analysed by non-linear regression using the Boltzmann equation y = LL + (UL – LL)/(1 + exp(T_m_ – x)/a) where LL and UL are the minimum and maximum fluorescence intensities, respectively (31).

#### In vitro transcription

pUC57-globin-Fluc was linearised with FspI and pUC57-ZIKV-Fluc was linearised with HindIII. RNA was transcribed using recombinant T7 polymerase at a final concentration of 50 ng/μL in transcription buffer (40 mM HEPES pH 7.5, 32 mM MgOAc, 40 mM DTT, 2 mM Spermidine, 10 mM NTPs, 0.2 U/μL RNaseOUT (Invitrogen)) for 2-4 hours at 37 °C. RNA was purified by DNaseI treatment, acidic phenol extraction and ethanol precipitation. Residual nucleotides were removed using Illustra MicroSpin G-50 columns (GE Healthcare). RNA was capped using ScriptCap system and ScriptCap 2′-*O*-methyltransferase (CellScript).

#### In vitro translation

IFIT proteins or complexes were diluted in bovine serum albumin (BSA) diluent buffer (0.5 mg/mL BSA, 20 mM Tris pH 7.5, 160 mM KCl, 5 % glycerol, 2 mM DTT, 1U/μL RNaseOUT), and incubated with 125 ng reporter RNA for 15 minutes at 37 °C to allow RNA binding. *In vitro* translation was performed using the Flexi Rabbit Reticulocyte Lysate (RRL) System (Promega) for 90 minutes at 30 °C. Reactions were terminated by incubation on ice, followed by addition of 50 volumes of passive lysis buffer (Promega). After addition of an equal volume of firefly luciferase substrate (20 mM Tricine, 2.67 mM MgSO_4_, 0.1 mM EDTA, 33.3 mM DTT, 530 μM ATP, 270 μM Acetyl-CoA, 30 mg/mL luciferin, 250 μM magnesium carbonate hydroxide), luciferase signal was measured by GloMax (Promega). Luciferase values were normalised to the diluent buffer-only control for each experiment.

#### Analysis of the IFIT1-mRNA interaction by inhibition of primer extension (toeprinting)

IFIT1/mRNA interaction was performed essentially as described previously (13) with minor modifications. Cap0-ZV or cap0-β-globin reporter mRNAs (1 nM) were incubated with IFIT1 or IFIT1 containing complexes (at concentrations indicated in Figures) for 10 min at 37 °C in 20 μL reactions containing 20 mM Tris pH 7.5, 100 mM KCl, 2.5 mM MgCl_2_, 1 mM ATP, 0.2 mM GTP, 1 mM DTT, 0.25 mM spermidine and 0.5 mg/mL BSA. IFIT1/mRNA interaction was monitored by inhibition of primer extension using avian myeloblastosis virus reverse transcriptase (2.5 U) (Promega) and a ^32^P-labelled primer in the presence of 4 mM MgCl_2_ and 0.5 mM dNTPs. Full length and truncated cDNA products were separated in a denaturing 6 % acrylamide gel and detected by autoradiography using an FLA7000 Typhoon Scanner (GE). Analysis was performed using Image-Quant TL.

#### Stable isotope labelling with amino acids in cell culture and immunoprecipitation

HEK293T cells were cultured in Arg/Lys-free DMEM, supplemented with light (R0K0), medium (R6K4) or heavy (R10K8) amino acids, as described (32). 1x10^7^ cells were transfected with 10 μg plasmid DNA using Lipofectamine 2000 (ThermoFisher). After 24 hours, media was replaced to contain 1000 U/mL human interferon α-2a (Roferon-A, Roche) for a further 16 hours. Cells were harvested in lysis buffer (50 mM Tris pH 7.5, 150 mM NaCl, 1 mM EDTA, 1 % Triton X100) containing 1:200 Protease Inhibitor Cocktail Set III (Merck) and 1:200 Benzonase nuclease (Sigma-Aldrich). Lysates were normalised to 3 mg/mL of protein before incubation with anti-FLAG-M2 affinity gel (Sigma) at 4 °C for 18 hours. Beads were washed 3 times in Tris-buffered saline, then resuspended in 2X SDS-sample buffer (Invitrogen) and boiled for 5 minutes to elute bound proteins.

#### LC-MS/MS and sample preparation

Following immunoprecipitation, the combined samples were subjected to SDS-PAGE electrophoresis on a precast gel, and extracted as a single band for in-gel trypsinisation. The resulting peptides were fractionated using an Ultimate 3000 nanoHPLC system in line with an Orbitrap Fusion Tribrid mass spectrometer (Thermo Scientific). In brief, peptides in 1% (vol/vol) formic acid were injected onto an Acclaim PepMap C18 nano-trap column (Thermo Scientific). After washing with 0.5 % (vol/vol) acetonitrile 0.1 % (vol/vol) formic acid peptides were resolved on a 250 mm × 75 μm Acclaim PepMap C18 reverse phase analytical column (Thermo Scientific) over a 150 min organic gradient, using 7 gradient segments (1-6 % solvent B over 1 min., 6-15 % B over 58 min., 15-32 %B over 58 min., 32-40 %B over 5 min., 40-90 %B over 1 min., held at 90 %B for 6 min and then reduced to 1 %B over 1 min.) with a flow rate of 300 nL min^−1^. Solvent A was 0.1 % formic acid and Solvent B was aqueous 80 % acetonitrile in 0.1 % formic acid. Peptides were ionized by nano-electrospray ionization at 2.0 kV using a stainless steel emitter with an internal diameter of 30 μm (Thermo Scientific) and a capillary temperature of 275 °C.

All spectra were acquired using an Orbitrap Fusion Tribrid mass spectrometer controlled by Xcalibur 2.1 software (Thermo Scientific) and operated in data-dependent acquisition mode. FTMS1 spectra were collected at a resolution of 120 000 over a scan range (m/z) of 350-1550, with an automatic gain control (AGC) target of 300 000 and a max injection time of 100 ms. Precursors were filtered using an Intensity Range of 1E4 to 1E20 and according to charge state (to include charge states 2-6) and with monoisotopic precursor selection. Previously interrogated precursors were excluded using a dynamic window (40 s +/-10 ppm). The MS2 precursors were isolated with a quadrupole mass filter set to a width of 1.4 m/z. ITMS2 spectra were collected with an AGC target of 20 000, max injection time of 40 ms and CID collision energy of 35 %.

#### Mass spectrometry data analysis

The raw data files were processed and quantified using MaxQuant v1.5.7.4 (33) and searched against the Uniprot Human database (70,550 entries, dated 19th September 2016) using the built-in Andromeda search engine. Peptide precursor mass tolerance was set a 4.5 ppm, and MS/MS tolerance was set at 0.5 Da. Search criteria included carbaminomethylation of Cys as a fixed modification. Oxidation of Met and N-terminal acetylation were selected as variable modifications. Quantification was based on Light (Arg 0, Lys 0), Medium (Arg 6, Lys 4), and Heavy (Arg 10, Lys 8) SILAC labels. Searches were performed with tryptic digestion, a minimum peptide length of seven amino acids, and a maximum of two missed cleavages were allowed. The reverse database search option was enabled and the maximum false discovery rate for both peptide and protein identifications was set to 0.01. Quantitation was performed using a mass precision of 2 ppm. The full MaxQuant output is provided as part of PRIDE submission PXD007584 permitting viewing of annotated spectra in MaxQuant v1.5.7.4. Downstream analysis was accomplished in the Perseus software (34). Contaminants and reverse database hits were removed, and protein ratios were log2-transformed. Proteins were considered to represent putative interaction partners if they showed a significant (t-test, p<0.05) increase in their abundance compared with the control pulldown and had to have been identified in at least two of the three replicates.

#### Protein structure modelling and analysis

Protein structure analysis and generation of protein structure images was performed using PyMOL (The PyMOL Molecular Graphics System, Version 2.0 Schrödinger, LLC). The IFIT3 model was generated by submitting the IFIT3 amino acid sequence to the Swiss Model server. The crystal structure of the IFIT2 domain-swapped homodimer (PDB:4G1T) was used as a template for model generation. The electrostatic surface potential of IFIT2 and the IFIT3 model were analysed using PDB2PQR and APBS software (35–37).

## Results

### IFIT1 co-precipitates IFIT2 and IFIT3 independently of RNA association

IFIT1 was originally identified as an RNA binding protein after being precipitated by 5’-ppp RNA from lysates of IFN stimulated HEK293 cells. IFIT2 and IFIT3, as well as several well-characterised RNA binding proteins, co-precipitated with IFIT1 (12). We used SILAC proteomics to examine if IFIT1 could interact directly with IFIT2 and IFIT3 in nuclease treated lysates from IFN-stimulated cells. HEK293 cells were passaged in differentially isotopically labelled media and transfected with either a plasmid expressing FLAG-tagged IFIT1, FLAG-tagged IFIT5 (previously reported not to interact with other IFITs (12, 29)), or an empty vector control. 24 hours after transfection the cells were treated with IFN-αand incubated for a further 16 hours. Preparation of nuclease treated cell lysates and pull-down experiments are described in Materials and Methods. Consistent with previous reports (38), IFIT1 was poorly overexpressed while IFIT5 was strongly expressed (Figure S1). However, as shown in Figure 1A, IFIT2 and IFIT3 co-precipitated with FLAG-tagged IFIT1, while FLAG-tagged IFIT5 did not precipitate other IFIT family members (Figure 1B). The full SILAC data set is available on the PRIDE server. These results independently confirm the interaction of IFIT1, 2 and 3 in IFN-stimulated cell lysates and further demonstrate that this interaction is maintained after nuclease treatment. Both IFIT2 and IFIT3 were enriched to a similar extent in the IFIT1 pull downs (Figure 1A).

**Figure 1.**
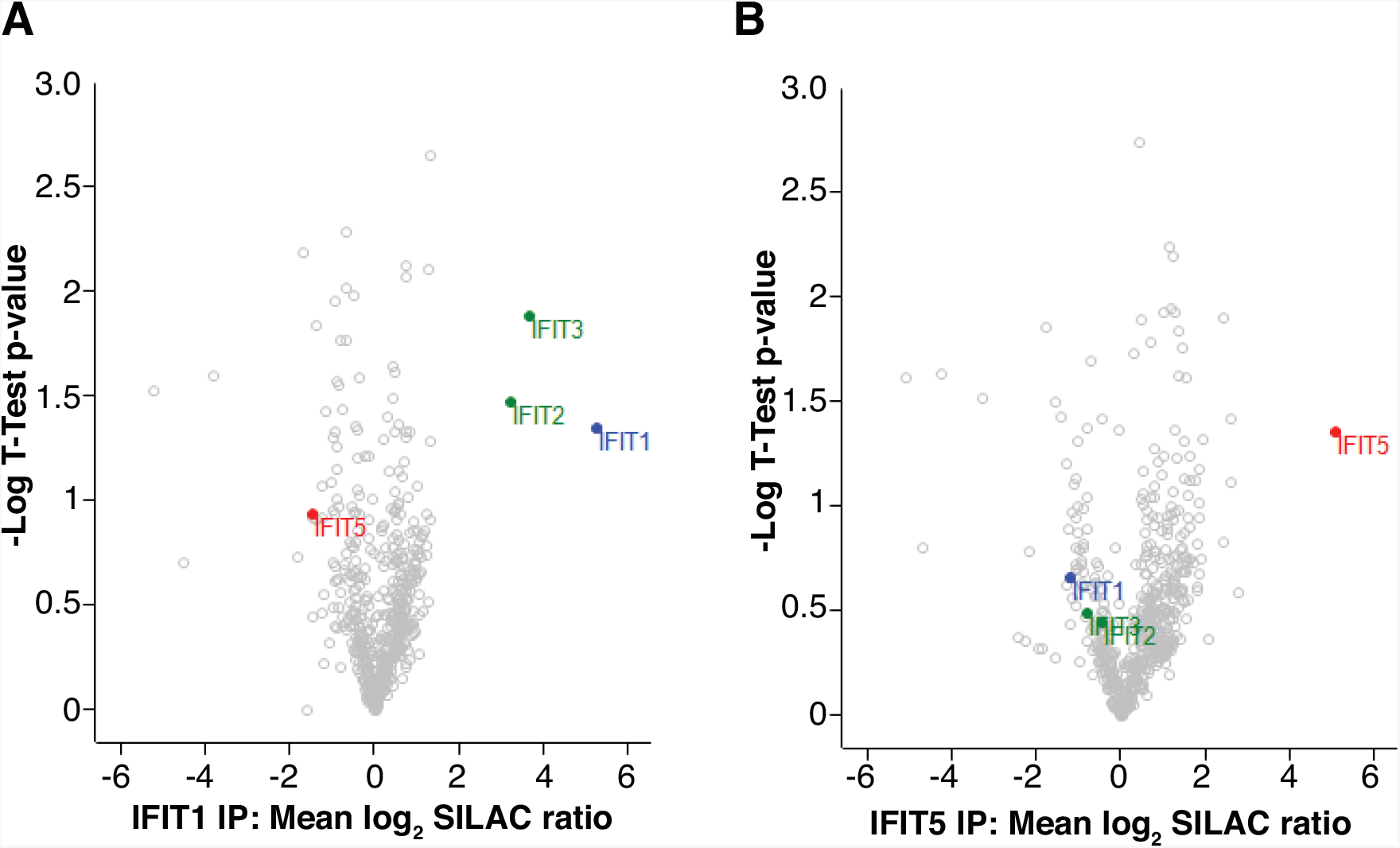
Interaction between IFIT1, IFIT2 and IFIT3 occurs independently of bound RNA. Volcano plots of SILAC proteomics data showing statistical analysis of proteins that immuno-precipitated with (A) FLAG-tagged IFIT1 or (B) FLAG-tagged IFIT5 following nuclease treatment of IFN-stimulated cell lysates. Data points corresponding to IFIT family members are coloured and labelled.

### IFIT1:IFIT2:IFIT3 hetero-complexes can be reconstituted from bacterially expressed proteins

To investigate human IFIT family oligomerisation and examine the influence of interaction with IFIT2 and IFIT3 on IFIT1 mRNA cap0 binding activity we reconstituted the IFIT1:IFIT2:IFIT3 hetero-complex *in vitro*. To this end, His-tagged IFIT1, IFIT2 and IFIT3 were individually expressed in bacteria and purified as described in Materials and Methods. The His-tag was removed from IFIT2 and IFIT3 but retained on IFIT1 for later detection. Each protein was subjected to size exclusion chromatography with multi angle light scattering (SEC-MALS) to analyse IFIT assembly. SEC-MALS reveals the molecular mass of species eluting at different volumes from a size exclusion column providing information about the oligomeric state of these species. Varying concentrations of IFIT were analysed by SEC-MALS and, consistent with a recent report (BioRxiv: https://doi.org/10.1101/152850), IFIT1 oligomerised in a concentration dependent manner (Figure S2A). In contrast, IFIT2 eluted as two species corresponding to a stable dimer or tetramer (Figure S2B). Surprisingly, the IFIT2 dimer had a similar elution volume to the lowest concentration of IFIT examined. This demonstrates the importance of using the SEC-MALS technique which directly determines the mass of particles in solution from their Rayleigh scattering, instead of relying on the elution volumes of molecular weight standards during SEC to infer oligomeric state. When analysed alone, IFIT3 also eluted as a mostly dimeric species, with a smaller peak corresponding to a monomer (Figure S2C).

We next examined the oligomeric status of complexes containing mixtures of the individually purified IFIT proteins. When incubated at 4 °C for 1 hour IFIT1 and IFIT3 formed a stable complex that eluted with a molecular mass of 221 kDa (Figure 2A, peak a). Analysis of the protein in peak a by SDS-PAGE (gel inset) shows that IFIT1 and IFIT3 are equimolar, indicating that this complex represents stable IFIT1:IFIT3 tetramers. A later eluting species likely corresponding to IFIT1:IFIT3 dimers (123 kDa) is also evident (Figure 2A, peak b). IFIT1 and IFIT2 were incubated at 30 °C for 1 hour and analysed by SEC-MALS. The IFIT1:IFIT2 complex was less defined and eluted as multiple species ranging from 148 to >234 kDa as measured by SEC-MALS (Figure S2D). While the most abundant IFIT1:IFIT2 complex form is likely tetrameric, the presence of several overlapping peaks precludes reliable determination of specific molecular mass and hence oligomeric state.

**Figure 2.**
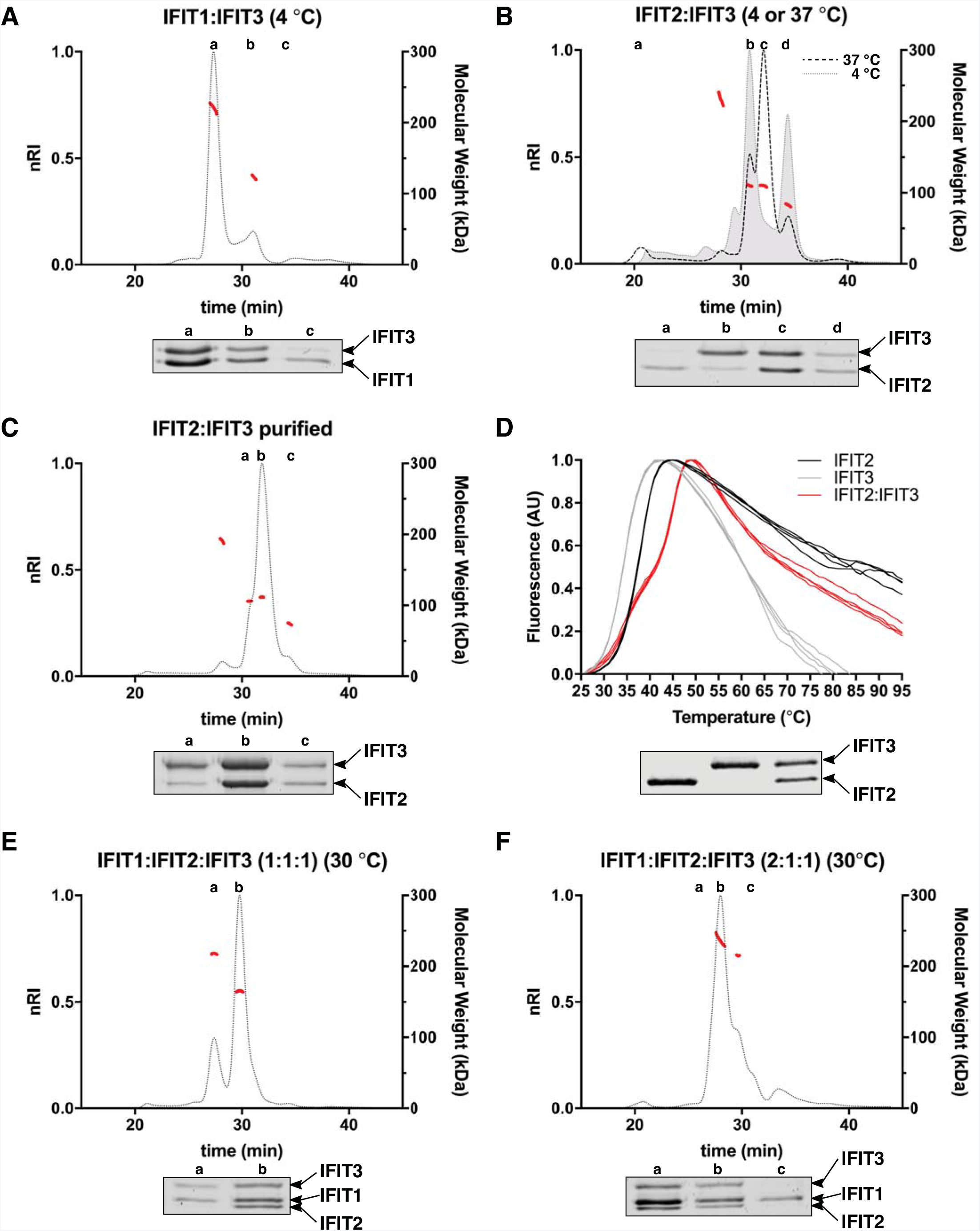
*In vitro* reconstitution of IFIT oligomeric assemblies. (**A,B**) Indicated IFIT complexes were assembled and analysed by SEC-MALS. (**C**) SEC-MALS analysis of the purified IFIT2:IFIT3 dimer alone or after incubation with an equimolar amount (**E**) or a 2-fold excess (**F**) of IFIT1. (**D**) Differential scanning fluorimetry analysis of IFIT2, IFIT3 and IFIT2:IFIT3 dimers. Gel inset shows SDS-PAGE analysis of proteins used for analysis. For A-C, D and F, complexes were formed at the indicated temperatures. Normalised differential refractive index (nRI) is shown as dotted or broken lines on the left *y-axis*. Calculated molecular masses (kDa) of eluting species are shown as solid, red lines on the right *y-axis*. Gel insets below each trace show SDS-PAGE analysis of each run. Protein gel lanes and corresponding peaks are indicated by lower case letters. The position of IFIT1, IFIT2 and IFIT3 on the protein gels is indicated. The calculated molecular masses of individual IFITs are: IFIT1(with 6His tag)-58.6 kDa, IFIT2-54.9 kD and IFIT3-56.2 kDa.

Because IFIT2 and IFIT3 were enriched to a similar degree in our IFIT1 pull down SILAC experiments we hypothesised that IFIT1 interacts with a heterodimer of IFIT2 and IFIT3. We therefore next examined if IFIT2 and IFIT3 could interact. Equal amounts of IFIT2 and IFIT3 were mixed and incubated at 4 °C for 1 hour. When analysed by SEC-MALS two peaks corresponding to IFIT2 and IFIT3 homodimers were clearly separated (Figure 2B, grey dotted line). In contrast, when IFIT2 and IFIT3 were instead incubated at 37 °C for 1 hour an IFIT2:IFIT3 heterodimer was formed (Figure 2B, black dashed line). The molecular weight of the eluting species (110 kDa) and analysis by SDS-PAGE (gel inset) are consistent with this complex representing an IFIT2:IFIT3 heterodimer. To test the stability of the IFIT2:IFIT3 heterodimer, corresponding peak fractions were concentrated and reanalysed by SEC-MALS. As evident in Figure 2C, there was no observed dissociation of the complex into constituent components, confirming that the IFIT2:IFIT3 complex remains stable.

To determine the relative stabilities of the different dimeric complexes, we examined IFIT2, IFIT3 and the IFIT2:IFIT3 heterodimer using differential scanning fluorimetry. Using this approach, protein unfolding is monitored by measuring the signal from a dye that fluoresces in hydrophobic environments, at increasing temperature. More stable proteins unfold at higher temperatures than less stable proteins. Unfolding exposes hydrophobic regions of the protein making them more accessible to the dye. Figure 2D shows the thermal melt curves for IFIT2 and IFIT3 homodimers and the IFIT2:IFIT3 heterodimer. The protein used in this assay is shown in the gel inset below the graph. IFIT2 and IFIT3 exhibit a monophasic melt curve from with T_m_’s of 37.3 °C and 34.1 °C, respectively, were calculated as described in the Material and Methods. This indicates that the IFIT2 is more thermodynamically stable than IFIT3. In contrast, the IFIT2:IFIT3 heterodimer exhibits a biphasic melt curve from which a reliable T_m_ could not be calculated. However, as is evident from comparison with the IFIT2 and IFIT3 melt curves that the melting temperature of the IFIT2:IFIT3 heterodimer, and thus the stability, is greater than that of either the homodimers alone.

Having successfully formed a stable IFIT2:IFIT3 complex, we attempted to reconstitute the full IFIT1:IFIT2:IFIT3 heterotrimeric complex. An equimolar ratio of IFIT1 was incubated with the purified (preformed) IFIT2:IFIT3 heterodimeric complex. Most of the IFIT1 was incorporated into a larger complex that eluted with a molecular mass of 165 kDa corresponding to a trimer (Figure 2E). Analysis of this complex by SDS-PAGE (gel inset) reveals that it contains equimolar amounts of IFIT1, IFIT2 and IFIT3. The remainder of the IFIT1 complexed with free IFIT3, a minor contaminant of the purified IFIT2:IFIT3 heterodimer, and eluted slightly earlier as a hetero-tetrameric complex with a molecular mass of 218 kDa. Finally, we incubated the IFIT2:IFIT3 heterodimer with a two-fold molar excess of IFIT1 at 30 °C for 1 hour. Analysis by SEC-MALS reveals a clear shift of the heterotrimeric complex to a heterotetrameric complex with a molecular mass of 236 kDa (Figure 2F). SDS-PAGE analysis of this complex (gel inset) reveals a molar excess of IFIT1 over IFIT2 and IFIT3 indicating that the heterotetrameric complex consists of two IFIT1 molecules to one molecule each of IFIT2 and 3 (compare gel insets in Figures 2 E and F). The IFIT1:IFIT2:IFIT3 heterotetramer was less stable than the trimer, precipitating when incubated at 4 °C. Subsequently, only the IFIT1:IFIT2:IFIT3 heterotrimer was used for further experiments.

### IFIT3 and IFIT2:IFIT3 stimulate the cap0-RNA translation inhibition activity of IFIT1

To investigate the impact of hetero-oligomerisation on the ability of IFIT1 to inhibit cap0 dependent translation we used a similar approach to that previously described by Young *et al.* (22). *In vitro* transcribed mRNAs comprising a firefly luciferase (Fluc) reporter flanked by either the human β- globin or zika virus (ZV) 5' and 3' untranslated regions (UTRs) were post-transcriptionally capped as described in the Materials and Methods. The β-globin and ZV constructs represent mRNAs with weak or strong secondary structures in the 5' UTR, respectively. Schematics of the two constructs are shown in Figure 3A. These mRNAs were incubated with different IFIT heterocomplexes before addition of rabbit reticulocyte lysate (RRL). Translation was quantified by measuring the luminescence from the Fluc reporter. The amount of IFIT1 added was equalised by western blotting against the His-tag on the purified IFIT1 in each complex. The upper panel of Figure 3B shows an SDS-PAGE analysis of the protein complexes used in these experiments while the lower panel shows an anti-His-tag western blot of the same complexes. The linearity of the western blot signal to protein concentration was confirmed as shown in Figure S3.

**Figure 3.**
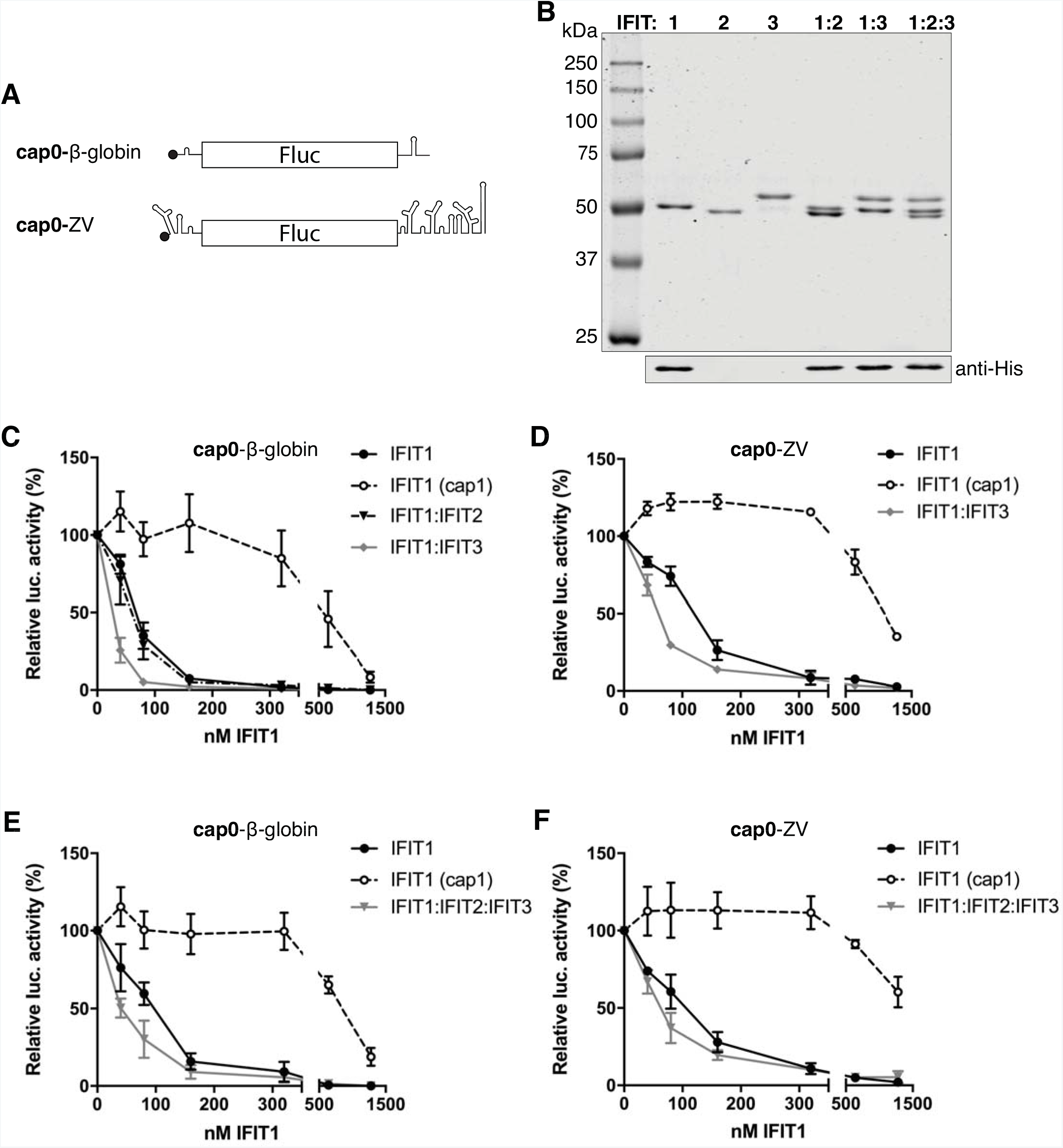
IFIT3 and IFIT2:IFIT3 enhance IFIT1 cap0 translation inhibition *in vitro*. (**A**)Schematic representation of the cap0-mRNAs used in *in vitro* translation assays in RRL. (**B**) IFIT1 containing complexes included in *in vitro* translation assays*Upper panel*, Coomassie stained SDS-PAGE gel analysis of individually purified and complexed IFITs. *Lower panel*, analysis of the same samples as in the upper panel by western blotting against the His-tag. Note, the His-tag was removed from IFIT2 and IFIT3 but not IFIT1 during purification. (**C-F**) Luciferase activity from RRL incubated with cap0- or cap1-β-globin Fluc RNA (C, E) or cap0- or cap1-ZV Fluc RNA (D, F) in the presence of increasing concentrations of IFIT1 or IFIT1 containing complexes as indicated. Data are normalised to the luciferase activity in the absence of IFITs and shown as the mean ± the standard error of three separate experiments.

IFIT1 inhibited translation of the cap0-β-globin Fluc reporter in a concentration-dependent manner (Figure 3C). Pre-incubation of IFIT1 and IFIT2 had no detectable impact on the inhibition of translation of the cap0-β-globin Fluc reporter and so was not examined further. In contrast, complexing with IFIT3 reproducibly decreased the concentration of IFIT1 required to cause 50 % inhibition of the same reporter (Figure 3C). IFIT1 inhibited translation of the cap0-ZV reporter less than the β-globin reporter overall, but complexing with IFIT3 again enhanced IFIT1’s translation inhibitory effect (Figure 3D). Because the heterotetrameric IFIT1:IFIT2:IFIT3 complex was unstable we focused on the heterotrimeric complex for further analysis. As can be seen in Figure 3E and 3F, stimulation of IFIT1 translation inhibition was reproducibly observed in the context of the IFIT1:IFIT2:IFIT3 complex. Reporter mRNAs bearing cap1 structures were also examined (Figure 3C-F and Figure S4). Inhibition of cap1 mRNAs was only observed at very high concentrations of IFIT1.

### IFIT3 and IFIT2:IFIT3 enhance cap0-RNA binding by IFIT1

We and others have previously demonstrated that IFIT1 binds preferentially to mRNA with a cap0 structure at the 5 end (13, 14). Using our purified complexes, we next examined if the interaction of IFIT1 with cap0 mRNA was altered when part of a larger IFIT1:IFIT3 or IFIT1:IFIT2:IFIT3 complex. We used a primer extension inhibition assay to monitor the IFIT1/cap0 mRNA interaction as previously described (13). An advantage of this technique over other methods to analyse protein-RNA interactions, such as electrophoretic mobility shift assays, is that the primer extension reaction is performed in equilibrium binding conditions. IFIT1 alone or as part of an IFIT heterocomplex was incubated with the *in vitro* transcribed and capped model β-globin and ZV mRNAs prior to the addition of a radiolabelled primer that binds within the Fluc mRNA sequence. A reverse transcription reaction is performed in which a full-length cDNA is produced in the absence of IFIT1, whereas a 7 nucleotide truncated cDNA, corresponding to the length of the IFIT1 RNA-binding surface, is produced in the presence of IFIT1 (13). The cDNA products are subsequently separated by denaturing PAGE and detected by autoradiography. IFIT protein complexes shown in Figure 3B were used in the binding reactions. Representative autoradiographs are shown in Figure 4A. Quantification of the cDNA products was performed as described in the Materials and Methods and the binding curves shown in Figure 4B and C show the fraction of RNA bound at varying IFIT concentrations.

**Figure 4.**
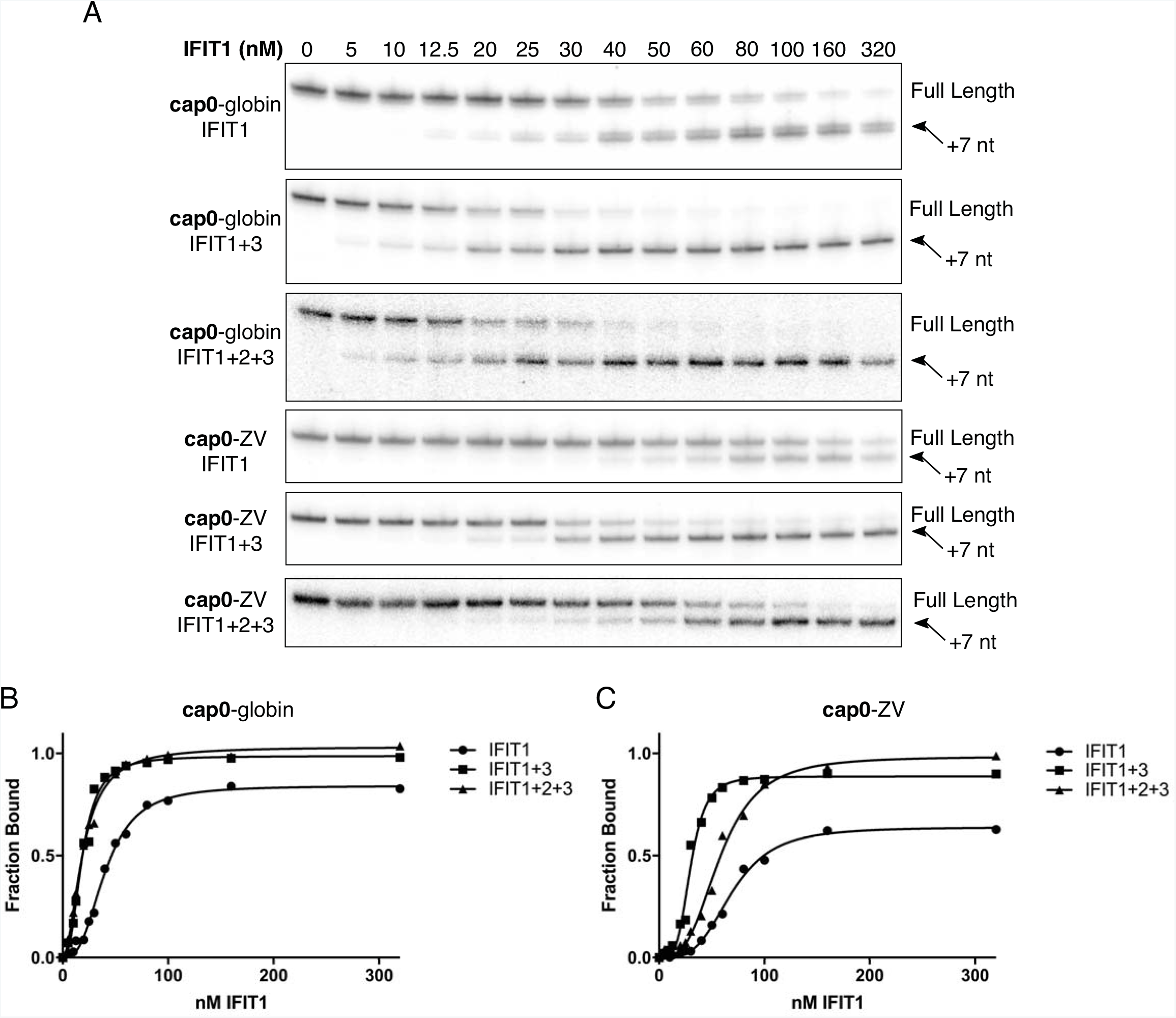
IFIT3 containing complexes stabilise IFIT1 binding to cap0 RNA. (**A**)Toeprinting analysis of the interaction of IFIT1 and IFIT1 containing complexes with cap0 RNA. The full-length and 7 nucleotide (nt) truncated cDNA product produced by IFIT1 binding are indicated. Protein complexes and RNAs are the same as those used in Figure 3. (B, C) Graphs represent fraction of RNA bound by IFIT1 and IFIT1 containing complexes at varying IFIT1 concentrations. Curves representative of three separate experiments were fitted using the nonlinear Hill equation, Fraction^[bound]^=[IFIT1]^h^ • Fraction^[bound]^_max_/([IFIT1]^h^+K^h^_1/2,app_) from data where [IFIT1] was ≥ 10 • [mRNA]. K_1/2, *_app_*_and Hill coefficients (*h*) are listed in the text.

As previously reported (13), IFIT1 binds cap0-β-globin mRNA with very high affinity (K_1/2,app_= 40 ± 1.2 nM, *h* = 2.8 ± 0.2). Complexing with IFIT3 or the IFIT2:IFIT3 heterodimer reduces the K_1/2,app_ of IFIT1 on this mRNA to 19 ± 0.8 nM ( *h* = 2.5 ± 0.2) and 19 ± 0.7 nM ( *h* = 1.98 ± 0.1), respectively. IFIT1 heterocomplexes also has the effect of saturating the binding on the mRNA as evident in the autoradiograms. This effect was even more pronounced when the cap0-ZV reporter was analysed. On this more structured RNA the K_1/2,app_ of IFIT1 binding was 69 ± 1.8 nM ( *h* = 3.5 ± 0.3) and only reached 60% saturation. In contrast, when complexed with IFIT3 or the IFIT2:IFIT3 heterodimer IFIT1 bound the ZV reporter with a K_1/2,app_ of 29 ± 1.1 nM ( *h* = 4.1 ± 0.6) and 58 ± 2.3 nM ( *h* = 3.1 ± 0.32), respectively. Again, as is clear from the autoradiograms, addition of IFIT3 or the IFIT2:IFIT3 heterodimer led to saturation of RNA binding. In all cases, and similar to our previous findings (13), the Hill coefficient was greater than 1, indicating a degree of cooperativity in IFIT1 cap0-mRNA binding.

### IFIT1 and IFIT3 interact through a C-terminal motif

Our data reveal that IFIT3 or the IFIT2:IFIT3 heterodimer can stimulate cap0 mRNA binding, and thus translation inhibition, by IFIT1. We next sought to identify how IFIT1 and IFIT3 interact. Murine Ifit3, which does not precipitate with murine Ifit1b1 (14), has a large deletion at the C terminus (Figure S5) when compared to human IFIT3. As a result of this deletion, mouse Ifit3 lacks a ‘YxxxL’ structural motif present in both human IFIT3 and IFIT1 (Figure 5A and B) recently reported to promote IFIT1 concentration dependent dimerisation (BioRxiv: https://doi.org/10.1101/152850). Our SEC-MALS analysis demonstrates that the IFIT1:IFIT3 interaction is more stable than the IFIT1:IFIT1 interaction (compare Figure 2A and Figure S2A). We therefore hypothesised that the proposed IFIT1 dimerisation motif is the site of interaction between IFIT1 and IFIT3.

**Figure 5.**
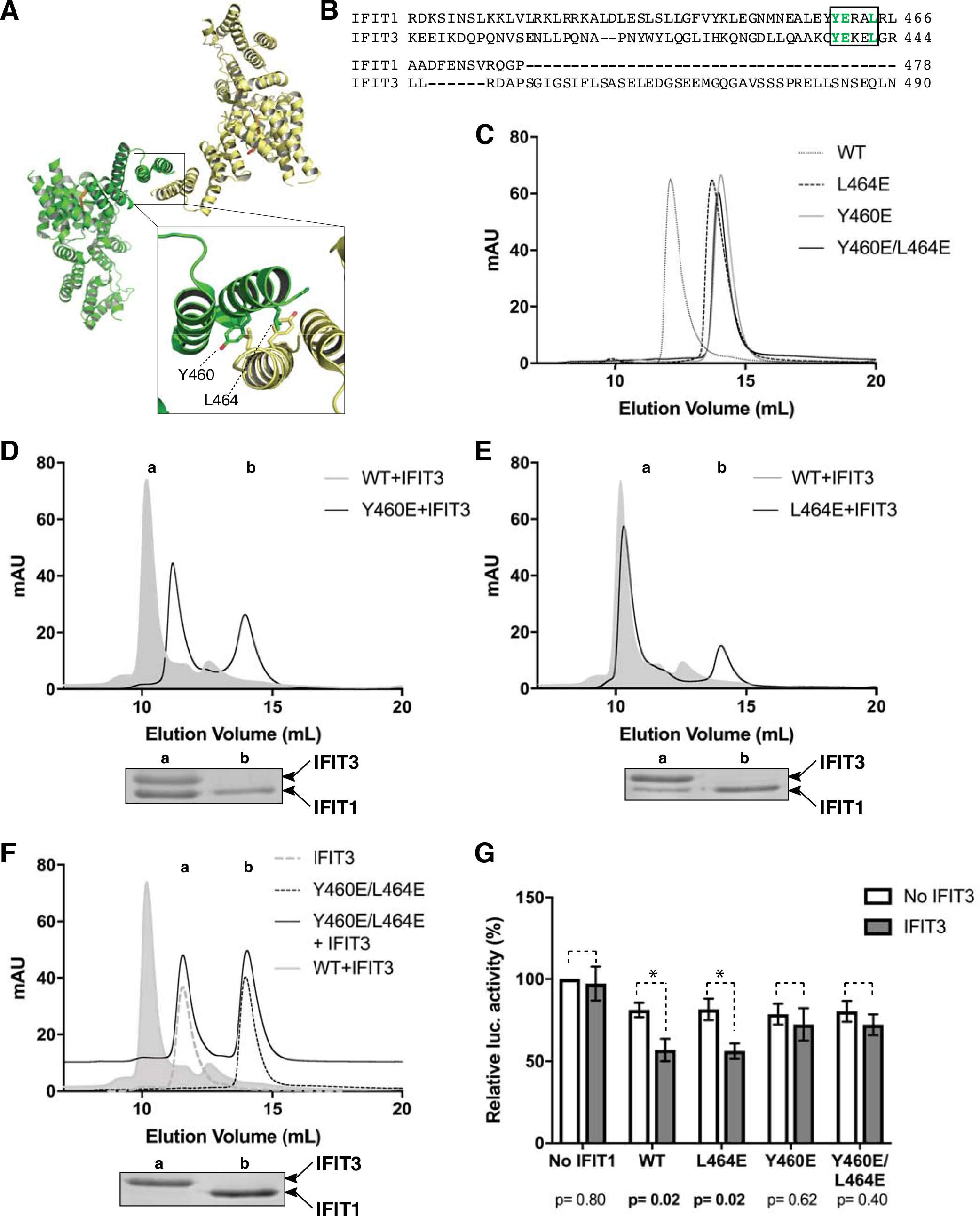
Mutational analysis reveals a key IFIT1:IFIT3 interaction motif. (**A**)Crystal structure of wild type IFIT1 (PDB: 5W5H). The two chains in the asymmetric unit are coloured yellow and green with m7GpppAAAA bound in the cap-binding pocket highlighted in orange. The interface region between the two IFIT1 molecules with Y460 and L464 side chains is enlarged (only labelled for the chain coloured green). (**B**) Sequence alignment of the C-terminal regions of IFIT1 and IFIT3 generated by Clustal Omega. The ‘YxxxL’ motif is boxed. (**C**, **D, E** and **F**) UV_280_ absorbance traces of SEC analysis (SuperdexS200 Increase 10/300 column) of wild type (WT) and mutant IFIT1 alone or incubated with IFIT3. Gel insets below each trace show SDS-PAGE analysis of each run. ( **D, E** and **F**) Protein gel lanes and corresponding peaks are indicated by lower case letters. The position of IFIT1 and IFIT3 on the protein gels is indicated. The Y460E/L464E+IFIT3 trace IFIT1+IFIT3 is shown in grey shadow for reference. (**G**) Luciferase activity from RRL incubated with cap0-β-globin Fluc RNA and WT or mutant IFIT1 with or without IFIT3 as indicated. Data are normalised to the luciferase activity in the absence of IFIT1 and shown as the mean ± the standard error of three separate experiments. Statistical analysis was performed comparing conditions joined by dotted lines using an unpaired, two-tailed Students T-test. P values are indicated, and * denotes statistical significance.

We generated a panel of IFIT1 mutants based on the IFIT1 dimer crystal structure (PDB: 5W5H). The point mutants Y460E and L464E expressed and purified similarly to the wild type protein. Both mutants eluted as monomeric species during SEC (Figure 5C) consistent with disruption of IFIT1 homodimerisation as previously reported (BioRxiv: https://doi.org/10.1101/152850). However, the L464E substitution only had a modest effect on the interaction of IFIT1 with IFIT3 (Figure 5D). While a small portion of IFIT1-L464E eluted as a monomer, a stable tetramer with IFIT3 was the predominant species, like the wild type IFIT1 shown in Figure 2A. The Y460E substitution destabilises IFIT1:IFIT3 oligomerisation to a greater extent than the L464E substitution, with less IFIT1-Y460E sequestered to the IFIT1:IFIT3 complex (Figure 5E) but, as is clear in the SDS-PAGE analysis of this experiment (gel inset), IFIT1-Y460E can still associate with IFIT3. We therefore generated a Y460E/L464E double mutant of IFIT1 (IFIT1-YL). IFIT1-YL eluted as a monomeric species on SEC, similarly to the Y460E and L464E single mutants (Figure 5C). However, in contrast to these two single point mutants, IFIT1-YL did not interact with IFIT3 during SEC (Figure 5F).

Disruption of dimerisation was previously reported not to affect the translation inhibition activity of IFIT1 (BioRxiv: https://doi.org/10.1101/152850 and Figure 5G, white bars). Therefore, we examined what impact mutations in the ‘YxxxL’ motif had on the ability of IFIT3 to stimulate IFIT1 translation inhibition activity. IFIT1 mutants were combined with IFIT3, reporter mRNA and RRL as described in Materials and Methods and Fluc dependent luminescence was measured (Figure 5G). At 40 nM IFIT1, ‘YxxxL’ motif mutants displayed similar translation inhibition. IFIT3 significantly enhanced translation inhibition of both wild type IFIT1 and the IFIT1-L464E mutant. In contrast, IFIT3 did not stimulate translation inhibition of either the IFIT1-Y460E mutant or the L464E/Y460E double mutant.

## Discussion

The IFIT family of ISGs are among the most highly upregulated proteins during cellular response to viral infection and, while a role for IFITs in regulating translation has long been postulated, the mechanisms by which these proteins function are only recently being revealed. Here, we have presented, to our knowledge, the first *in vitro* reconstitution of the IFIT1:IFIT2:IFIT3 complex. This has enabled us to examine the impact of hetero oligomerisation on RNA recognition by IFIT1 and to begin to understand how this complex assembles.

### IFIT oligomerisation

We initially confirmed that IFIT1, IFIT2 and IFIT3 interact in lysates from HEK293T cells using proteomics. In contrast to previous studies (12, 14) we used nuclease treated samples confirming that this interaction occurs in cells in an RNA-independent manner. Moreover, we confirmed that IFIT1 could interact directly with IFIT2 or IFIT3 by SEC. We then employed SEC-MALS to examine IFIT heterocomplex assembly for the first time. Unlike standard SEC, when coupled with protein gel analysis SEC-MALS enables reliable identification of oligomeric status. Figure 6 shows schematic representations of each of the IFIT complexes reconstituted *in vitro* in this study. Consistent with previous reports ((11) and BioRxiv: https://doi.org/10.1101/152850), IFIT1 was found to reversibly dimerise in a concentration dependent manner (Figure S2A). While both IFIT2 and IFIT3 individually form stable homodimers as previously reported (13, 7), an unexpected tetrameric form of IFIT2 was also detected (Figure S2B and C). Surprisingly, we found that the IFIT1:IFIT3 interaction is more stable than the IFIT1:IFIT1 complex (compare Figures 2A and S2A). When analysed by SEC-MALS the predominant IFIT1:IFIT3 oligomeric state was a stable tetramer containing equimolar amounts of both proteins. The IFIT2 dimer eluted from the SEC at a volume expected for a monomeric species (Figure S2B) and as such had a similar elution volume as IFIT1. Very weak interaction of IFIT1 and IFIT2 was detected by SEC when these proteins were incubated at 4° C (*RF and TS unpublished data*).

**Figure 6.**
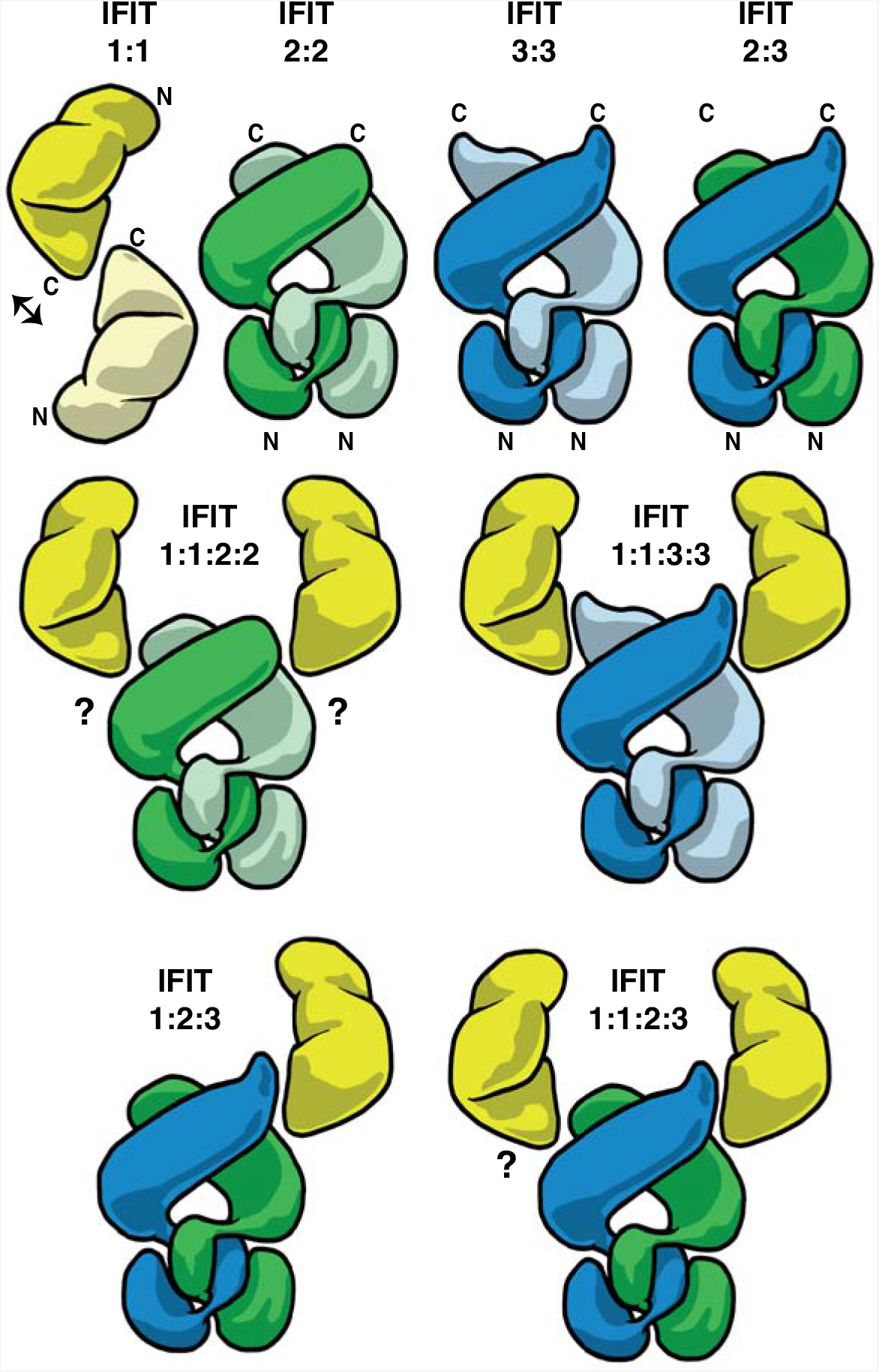
Schematic representation of the IFIT complexes analysed in this study. Cartoons depicting IFIT1 (yellow), IFIT2 (green) and IFIT3 (blue) complexes reconstituted *in vitro* from individually purified proteins. Weak interactions are indicated by reversible arrows. IFIT1 and IFIT2 structures and dimerisation interactions have been characterised by X-ray crystallography (BioRxiv: https://doi.org/10.1101/152850 and (30)). No experimentally derived structure for IFIT3 homo- or heterodimers are currently available. Therefore, IFIT3 and IFIT2:IFIT3 dimerisation, and IFIT1 heterooligomerisation are modelled on the IFIT1 dimer (PDB: 5W5H) and IFIT2 dimer (PDB: 4G1T) crystal structures and supported by experimental evidence as discussed in the text. The association of IFIT1 and IFIT2 is less well defined as indicated by the question marks.

However, when the incubation temperature was increased to 30 °C multiple oligomeric states of the IFIT1:IFIT2 complex were present (Figure S2D). These data suggest that IFIT1 interacts with IFIT2 in a different manner to IFIT3. Results of previous attempts to narrow down the IFIT1:IFIT2 binding interface using co-precipitation from lysates overexpressing IFIT2 truncation mutants were ambiguous and suggested a lack of specificity in the interaction (29).

Our SILAC data indicated that IFIT2 and IFIT3 were enriched to a similar degree in IFIT1-bound samples (Figure 1), suggesting that a heterocomplex of IFIT2:IFIT3 was the more likely form to interact with IFIT1 in the cell. This prompted us to investigate the IFIT2:IFIT3 interaction in more detail. The delayed elution of the dimeric IFIT2 from the size exclusion column fortuitously enabled us to analyse the IFIT2:IFIT3 interaction by SEC-MALS. Although no interaction between these two proteins was detected when incubated at 4 °C, we observed the appearance of a heterodimeric complex when these proteins were incubated at 37 °C (Figure 2B) indicating that addition of energy was required for this heterocomplex to form. The crystal structure of IFIT2 revealed that dimerisation of the protein was due to a domain swap of three α-helices that constitute one and a half TPRs in the N-terminal domain (7) (Figure S6A and shown schematically in Figure 6). Nucleic acid binding activity of IFIT2 was localised to the dimer interface surfaces of the C-terminal domain that form a large positively charged pocket (Figure S6B). The electrostatic surface potential of an IFIT3 molecular model, based on the IFIT2 crystal structure, is shown in Figure S6C. The IFIT3 model lacks a positively-charged nucleic acid binding surface like that of IFIT2.

The ability of IFIT3 to sequester IFIT2 into a heterodimeric form demonstrates that IFIT3 can disrupt the domain swapped architecture of IFIT2. It is therefore possible that IFIT3 could interact with IFIT2 in a similar domain swapped manner (modelled schematically in Figure 6, *top right*), consistent with TPRs 1-4 in the N terminus of IFIT2 being sufficient for the interaction as reported previously (29). Results of SEC analysis and differential scanning fluorimetry experiments indicate that the IFIT2:IFIT3 complex is more stable than separate IFIT2 and IFT3 homodimers (Figure 2C and D). The reason for this is currently unclear but comparison of the thermal melt curves of the three complexes shows that unlike the IFIT2 and IFIT3 homodimers, unfolding of the IFIT2:IFIT3 complex is biphasic (Figure 2D). A possible explanation for this is that the IFIT2 and IFIT3 proteins unfold uniformly, whereas the interacting regions of the IFIT2:IFIT3 heterocomplex are stabilised and unfold at different rates to the rest of the protein domains in the complex. The IFIT2:IFIT3 complex does not display unfolding kinetics averaged between those observed for IFIT2 and IFIT3 suggesting that the heterocomplex is indeed more stable than each of its constituent parts alone. High-resolution structures of the IFIT2:IFIT3 complex are required to confirm how these proteins interact and are a focus of our ongoing investigation.

Our IFIT2:IFIT3 interaction studies have important implications for our understanding of IFIT complex assembly and biology. Overexpression of IFIT2 was previously reported to induce apoptosis (29, 39), while co-expression with IFIT3 but not IFIT1 blocked this effect (29). Moreover, depletion of IFIT3 induced cell death in the U549 human carcinoma cell line, an effect potentiated by co-infection with Dengue virus (40). Our data would suggest that the homodimeric form of IFIT2 may be responsible for this phenotype, and that co-expression of IFIT3 may mitigate this effect by disrupting IFIT2 dimerisation. It is currently not clear why the cell would evolve such a mechanism for inducing programmed cell death. One potential hypothesis is that dysregulation of ISG induction, perhaps due to infection, could perturb the balance of IFIT2 and IFIT3, promoting cell death to restrict pathogen spread.

Combining IFIT1 in equimolar amounts or in two-fold excess with the purified IFIT2:IFIT3 heterodimer resulted in the formation of heterotrimeric or heterotetrameric IFIT complexes, respectively (Figure 2E and F and shown schematically in Figure 6, *bottom row*). Importantly, this complex closely resembles the IFIT1:IFIT2:IFIT3-containing 150-200 kDa complex previously identified in HeLa cell lysates (29), supporting its biological relevance. Unlike the IFIT1:IFIT3 tetramer, the IFIT1:IFIT2:IFIT3 tetramer was unstable and precipitated during incubation at 4 °C. In contrast, the heterotrimer was stable and enabled us to investigate the impact of IFIT2:IFIT3 complexing on the translation inhibition activity of IFIT1 (discussed below). A recent report described the crystal structure of wild type IFIT1 and revealed a dimerisation interface between C-terminal TPRs (BioRxiv: https://doi.org/10.1101/152850), responsible for the concentration dependent homo dimerisation of the protein, confirmed here. Importantly, mutation of this interface had no impact on the translation inhibition activity of IFIT1 (BioRxiv: https://doi.org/10.1101/152850 and Figure 5E) and Figure 5E). We observed that IFIT3 shares multiple key residues involved in the IFIT1 ‘YxxxL’ dimerisation motif, also in a C-terminal TPR. We therefore hypothesized that this motif is the site of interaction between IFIT1 and IFIT3. Mutational analysis of IFIT1 confirmed that this motif is indeed required for interaction with IFIT3 (Figure 5). The double-mutant IFIT1-Y460E/L464E did not interact with IFIT3 when analysed by SEC. Since the wild type IFIT1 homodimer is less stable than the IFIT1:IFIT3 complex, as shown by our SEC-MALS analysis, we propose that the functionally-important role of the ‘YxxxL’ motif in humans is mediating interaction between IFIT1 and IFIT3 (shown schematically in Figure 6). Importantly, the crystal structure of IFIT5 reveals that, although critical interacting residues may be conserved, they are buried in an interface with a terminal helix not present in IFIT1 (8–10), explaining why IFIT5 does not interact with IFIT3 ((12) and Figure 1). While the ‘YxxxL’ motif is conserved in mouse Ifit1b1, Ifit3 is truncated such that the ‘YxxxL’ motif is absent. In mice and other rodents, the Ifit1b1 gene has been duplicated (6) and it is possible that these extra IFITs could compensate for the disrupted murine Ifit1b1-Ifit3 interaction. This suggests species differences in the role of this motif, and in IFIT oligomerisation in general, that must be considered when examining phenotypes of small animal models used to examine the impact of IFIT depletion.

### Impact of oligomerisation on IFIT1 cap0 binding and translation inhibition

Our *in vitro* reconstitution of the human IFIT heterocomplex enabled us to examine the impact of oligomerisation on the cap0 mRNA binding and translation inhibition activity of IFIT1. We opted to use an RRL based translation inhibition assay system as this was previously used to demonstrate differential inhibitory effects of IFIT1 on mRNAs from parainfluenza virus 5 (22). We used two reporters, one with a well characterised (41), highly structured 5' UTR (ZV, *dG*=-33.4 kcal/mol) and one with very little structure (β-globin, *dG*=-10.5 kcal/mol), upstream of a Fluc reporter and with the corresponding 3' UTRs. IFIT1 strongly inhibited translation of both reporters (Figure 3), but had a stronger effect on the less-structured β-globin reporter, consistent with an emerging consensus that both the methylation state and RNA structure at the 5' end of an mRNA can influence its susceptibility to IFIT1 inhibition (9, 22, 23). IFIT2 had no detectable impact on the ability of IFIT1 to inhibit translation in the RRL system (Figure 3C). In contrast, IFIT3 and to a lesser extent the IFIT2:IFIT3 heterodimer enhanced the translation inhibition effect of IFIT1 on both reporters (Figure 3C-F).

It is possible that the RNA binding surface of IFIT1 may be extended by interaction with its binding partners. This has the potential to expand the repertoire of RNAs IFIT1 can interact with by providing additional stabilising interactions. Moreover, it is unclear why inclusion of IFIT3 alone has a stronger effect on translation inhibition than inclusion of the IFIT2:IFIT3 heterodimer. Since inclusion of IFIT2 alone had no impact on the translation inhibition activity of IFIT1 it is possible that the stimulatory effect is a result of the IFIT3 in the IFIT2:IFIT3 complex. The physiological role for this stimulation of binding is not immediately clear as we have previously demonstrated that IFIT1 binding to cap0-mRNA is sufficient to prevent association of the cap-binding translation initiation factor, eIF4F, *in vitro* (13). However, a requirement for co-expression of other IFITs for the full anti-viral activity of IFIT1 has been reported (12). During infection the enhanced binding of IFIT1, as a result of interaction with IFIT3, may promote IFIT1-dependent restriction in regions where access to cellular proteins is limited, such as in viral replication complexes.

While multiple cellular factors such as competing cap-binding proteins complicate analysis in cells, our *in vitro* mRNA binding assays reveal a clear cap0-mRNA binding advantage conferred by IFIT1 heterooligomerisation. When examined in our reverse transcriptase inhibition assays apparent IFIT1 binding was much less efficient for the structured cap0-ZV reporter than for the less structured β- globin mRNA, and in fact on the ZV construct failed to reach saturation (Figure 4). This is not due to incomplete capping of the mRNA as saturation is possible when IFIT3 is present. Instead, it is more likely that the reverse transcriptase may remove a proportion of the bound IFIT1 as it proceeds in a 5 to 3 direction resulting in the production of a full-length signal even if IFIT1 was initially bound. We conclude therefore that IFIT3, and the IFIT2:IFIT3 heterodimer, stabilise the interaction of IFIT1 with the cap0 mRNA in such a way that it is no longer removed by the reverse transcriptase. Interestingly, there was no evidence of a shift in the IFIT dependent toeprint to indicate that the other IFITs in the complex were interacting with the mRNA downstream of the IFIT1 cap0 binding cleft. IFIT5 changes conformation when transitioning between the apo- and RNA-bound state (8), positioning key residues for optimal RNA binding. The crystal structures of IFIT1 bound to different short RNAs show the protein is in a similar closed conformation as the RNA-bound structure of IFIT5 (9). It is therefore likely that IFIT1 cycles through a similar open/closed conformation to interact with RNA. Binding of IFIT3 may promote the closed conformation of IFIT1, stabilising its interaction with target RNAs. Future efforts will focus on obtaining high-resolution structures of IFIT1:IFIT3 and IFIT1:IFIT2:IFIT3 complexes with and without RNA to address these questions.

Together, our results demonstrate the first *in vitro* reconstitution of the human IFIT2:IFIT3 and IFIT1:IFIT2:IFIT3 heterocomplexes, provide novel details about IFIT complex assembly and reveal for the first time that IFIT3 enhances IFIT1 cap0-mRNA binding and translation inhibition. Our reconstituted complexes provide a solid foundation for future molecular analysis of IFIT assembly and function.

## Acknowledgements

We are grateful to Janet Deane for providing access to equipment for SEC-MALS experiments and to George Katibah and Kathleen Collins for the gift of the IFIT1 expression plasmid.

